# Physical-Chemical Features Selection Reveals That Differences in Dipeptide Compositions Correlate Most with Protein-Protein Interactions

**DOI:** 10.1101/2024.02.27.582345

**Authors:** Hamid Teimouri, Angela Medvedeva, Anatoly B. Kolomeisky

## Abstract

The ability to accurately predict protein-protein interactions is critically important for our understanding of major cellular processes. However, current experimental and computational approaches for identifying them are technically very challenging and still have limited success. We propose a new computational method for predicting protein-protein interactions using only primary sequence information. It utilizes a concept of physical-chemical similarity to determine which interactions will most probably occur. In our approach, the physical-chemical features of protein are extracted using bioinformatics tools for different organisms, and then they are utilized in a machine-learning method to identify successful protein-protein interactions via correlation analysis. It is found that the most important property that correlates most with the protein-protein interactions for all studied organisms is dipeptide amino acid compositions. The analysis is specifically applied to the bacterial two-component system that includes histidine kinase and transcriptional response regulators. Our theoretical approach provides a simple and robust method for quantifying the important details of complex mechanisms of biological processes.

## 1 Introduction

Protein-protein interactions (PPI), that can be viewed as the result of various biochemical reactions and electrostatic attractions, play a critical role in many cellular processes by supporting a variety of crucial biological functions. [22, 30] These functions range from signal transduction, such as stimulus-response coupling in bacteria [11, 25], and enzymatic regulations to the generation of immune responses [5, 31, 32]. Furthermore, protein-protein interactions are closely associated with the development and progress of various diseases, including viral pathogenesis [7], cancer [23], and neurodegenerative diseases [31, 36]. For example, neurological disorders such as Alzheimer’s disease, Parkinson’s disease, and Huntington’s disease, all have been linked to mutations that specifically disrupt PPIs that can prevent misfolding, leading to effectively irreversible aggregation of proteins [5].

Exact identification of PPIs in cellular systems remains a very difficult task. Several experimental techniques, including yeast two-hybrid (Y2H) screens [13, 21], mass spectroscopy [19, 33], and tandem affinity purification (TAP) [42, 54] have been developed in recent years for detecting them. However, despite some advances, determining PPIs in labs remains technically very challenging, time-consuming, and costly. Additionally, due to the complexity of underlying processes, these experimental methods often exhibit high rates of false positives and false negatives [52]. As a result, several computational methods have been proposed to assist in predicting protein interactions more accurately and efficiently [8, 12, 45]. Such theoretical methods not only support traditional wet lab experiments but also offer a more cost-effective means to quickly identify potentially interacting protein pairs across the huge space of the entire proteome [24]. Yet, the performance of most of these techniques declines when the supplemental additional biological information, like protein structure details, protein domains, or gene neighborhood information, are not available [24]. Hence, there is an immediate need to devise new computational strategies that could predict more reliably PPIs preferably relying only on the limited information coming from protein sequence data [44].

Because of the large volume of available biological information, in recent years, machine learning methods have emerged as powerful tools to complement traditional experimental techniques, enabling the analysis and prediction of PPIs from amino acid sequences [17, 20, 44]. However, many advanced machine-learning models, like deep neural networks, are black boxes, making it difficult to understand why they make specific predictions. Such methods do not provide insights into which features of the protein sequence are most relevant for these interactions. Moreover, traditional models frequently rely on simplistic representations of protein sequences, such as amino acid composition or a very limited set of physicochemical descriptors of proteins. On the other hand, despite the advancements achieved in the field of PPI prediction using machine learning, current approaches often overlook a crucial aspect – the specificity of protein-protein interactions within different biological systems. Biological processes are highly contextual, and protein interactions may vary significantly across diverse organisms and cellular environments. Existing machine-learning methods might not fully capture the species-specific patterns and nuances of the PPI networks, limiting their ability to provide robust predictions.

Here, we present a novel computational approach that addresses this crucial gap in the abilities of PPI prediction techniques. Our hypothesis is that interactions between different protein species correlate with their molecular properties. In this approach, we extract a comprehensive set of physicochemical features of proteins using a standard bioinformatic tool [6]. Then, the concept of physicochemical similarity between protein pairs is applied to identify which proteins might interact with each other. By incorporating species-specific features and training machine-learning models on organism-specific datasets, our method reveals the unique aspects of PPI networks in different organisms. We investigated six diverse datasets encompassing microorganisms, mammals, insects, and plants, allowing us to comprehensively capture the properties of PPI networks across different biological kingdoms. The protein-protein interaction prediction is modeled as a classification problem, applying the principles of supervised machine learning. By employing supervised machine-learning techniques, specifically Logistic Regression and Support Vector Machines (SVM), we demonstrate that a selected set of physicochemical protein features can effectively predict whether proteins will interact or not. Our analysis identifies that dipeptide compositions are universal factors across all studied organisms that best correlate with the possibility of PPIs. The proposed computational method provides an enhanced approach to understanding the characteristics of proteins associated with successful interactions.

## 2 Materials and Methods

### 2.1 Dataset and Data Pre-Processing

We considered protein-protein interactions in two types of living systems: 1) unicellular organisms, including bacteria *Escherichia coli* (EC2) and two distinct species of yeast including, *Saccharomyces cerevisiae* (SC5), and *Schizosaccharomyces pombe* (SP) ; and 2) multicellular organisms, including *Mus musculus* (MM), *Drosophila melanogaster* (DM2), and *Arabidopsis thaliana* (AT). The summary of all utilized information for different systems is presented in Table 1.

**Table 1.** Summary of protein-protein interactions datasets used in our computational study. Data obtained from Ref. [9].

| Label | Species | Proteins | PPI (positive/negative) |
| --- | --- | --- | --- |
| EC2 | Escherichia coli | 589 | 1167/1167 |
| SP | Schizosaccharomyces pombe | 904 | 742/742 |
| AT | Arabidopsis thaliana | 756 | 541/541 |
| MM | Mus musculus | 1088 | 500/500 |
| DM2 | Drosophila melanogaster | 658 | 321/321 |
| SC5 | Saccharomyces cerevisiae | 454 | 500/500 |

The data for each organism consisted of pairs of proteins and their corresponding sequences. We labeled each protein-protein pair in the dataset as 1 if they interact and 0 if they do not interact. For each organism, there was an equal number of protein-protein pairs that interact vs those that do not interact, as illustrated in Table 1. This allows us to minimize the bias in the analysis of data.

### 2.2 Generation of Physicochemical Descriptors for Proteins

From the amino acid sequence of each protein, we extracted a comprehensive set of physicochemical descriptors using the *propy* package [6]. The features were broadly classified into different categories, including charge, residue compositions (e.g., dipeptide composition), autocorrelations, chemical compositions, and sequence order features. Proteins containing non-natural amino acids were excluded from our dataset, as the *propy* package only identifies natural amino acids and we are also interested in finding PPIs only in real cellular systems.

For each protein, the quantitative values of the physicochemical properties have different numerical values. It is important to initially rescale all these values to fall between 0 and 1 so that every property is considered with a similar weight. To normalize this quantity to be in the range 0 and 1, we use the following rescaling expression,

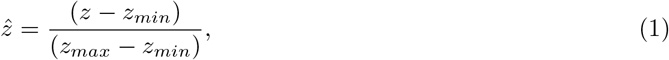

where *z* is the original value of the physicochemical property, *z*_*min*_ and *z*_*max*_ are limiting values for this property for all considered proteins, and 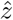 is the normalized one that is specifically utilized in the analysis.

### 2.3 Protein-Protein Interaction as a Classification Problem

By extracting various physicochemical features, we can mathematically represent each protein as a vector in a high-dimensional space of these properties. Let us consider two arbitrary proteins *A* and *B* for which there are *N* available physicochemical features. Their vector representations are *A* = [*A*_1_, *A*_2_, …, *A*_*N*_] and *B* = [*B*_1_, *B*_2_, …, *B*_*N*_], respectively. Thus, the difference between two vectors is given as another vector,

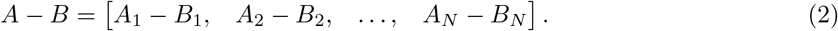

The process of identifying protein-protein interactions can be viewed as a supervised machine-learning problem. In our dataset, we assign an index *y*_*i*_ to each protein-protein pair. If the two proteins interact, *y*_*i*_ = 1, and if they do not interact, *y*_*i*_ = 0. The feature vector (with total *n* properties) **x**_*i*_ = {x_*i,1*_, x_*i,2*_, …, x_*i,n*_} for each protein-protein pair *i* describes differences of the two proteins in terms of individual features. Then, the Support Vector Machine (SVM) classification method [37] is employed for predicting protein-protein interactions from the differences in the physicochemical properties.

However, using all features leads to overfitting, as there is a significant gap between model performance on training data and test data. One needs to select a few most important properties to avoid overfitting, and the details are explained below.

### 2.4 Feature Selection Process

The number of possible physicochemical descriptors is very large, and many of these properties strongly correlate with each other. In such high-dimensional feature space, it is beneficial to identify a small subset of the most predictive features. This can be achieved mathematically by assigning zero weights to irrelevant or redundant features in regression and SVM methods. LASSO (The Least Absolute Shrinkage and Selection Operator) regression and Support Vector Machine are two prevalent techniques employed for shrinkage and feature selection [50]. The overall scheme for our procedure is presented Fig. 1.

**Figure 1.**
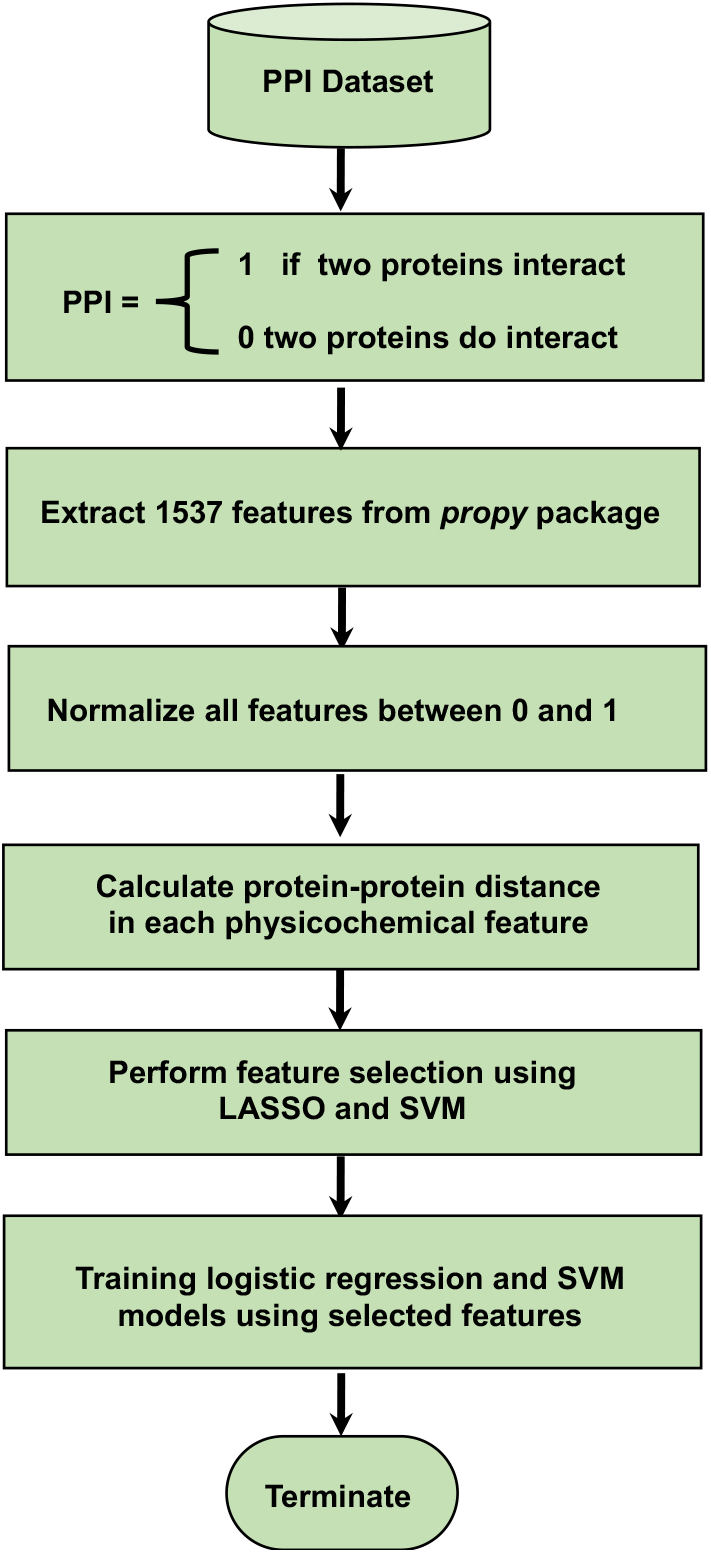
A flowchart of the method for selecting specific features for predicting protein-protein interactions.

The summaries of the LASSO and SVM feature selection methods are described in detail in Algorithm 1 and Algorithm 2, respectively.

#### 2.4.1 Evaluating Performance of Machine-Learning Models

In the evaluation of machine learning models, several metrics are commonly used to measure the performance of these models. Each of these metrics has its strengths and weaknesses. *Accuracy*, which is one of the most intuitive metrics, represents the proportion of correctly classified instances (both true positives and true negatives) to the total number of instances. This quantity can be evaluated via

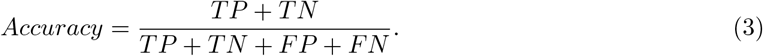

##### Algorithm 1

Feature Selection Using the Lasso Method

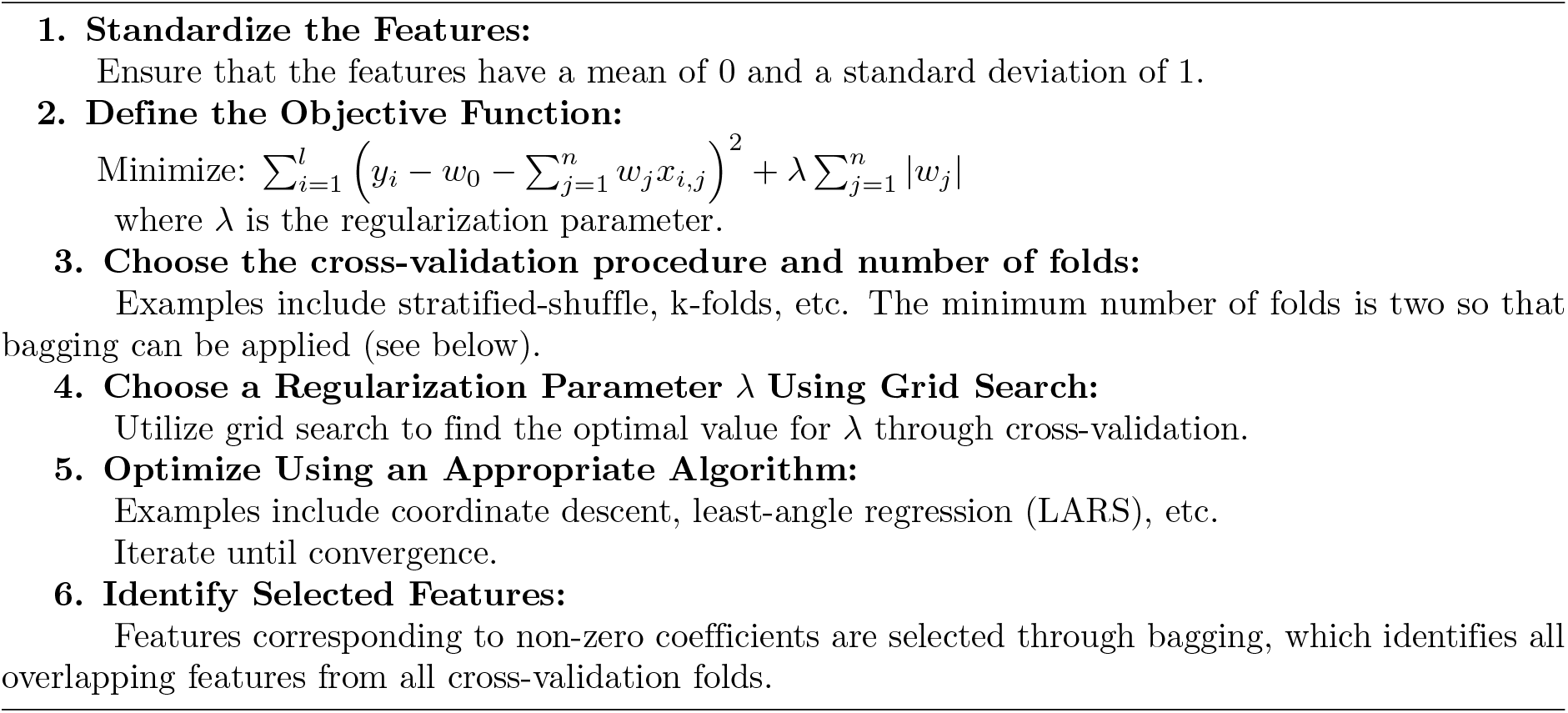

##### Algorithm 2

Feature Selection Using the SVM Method

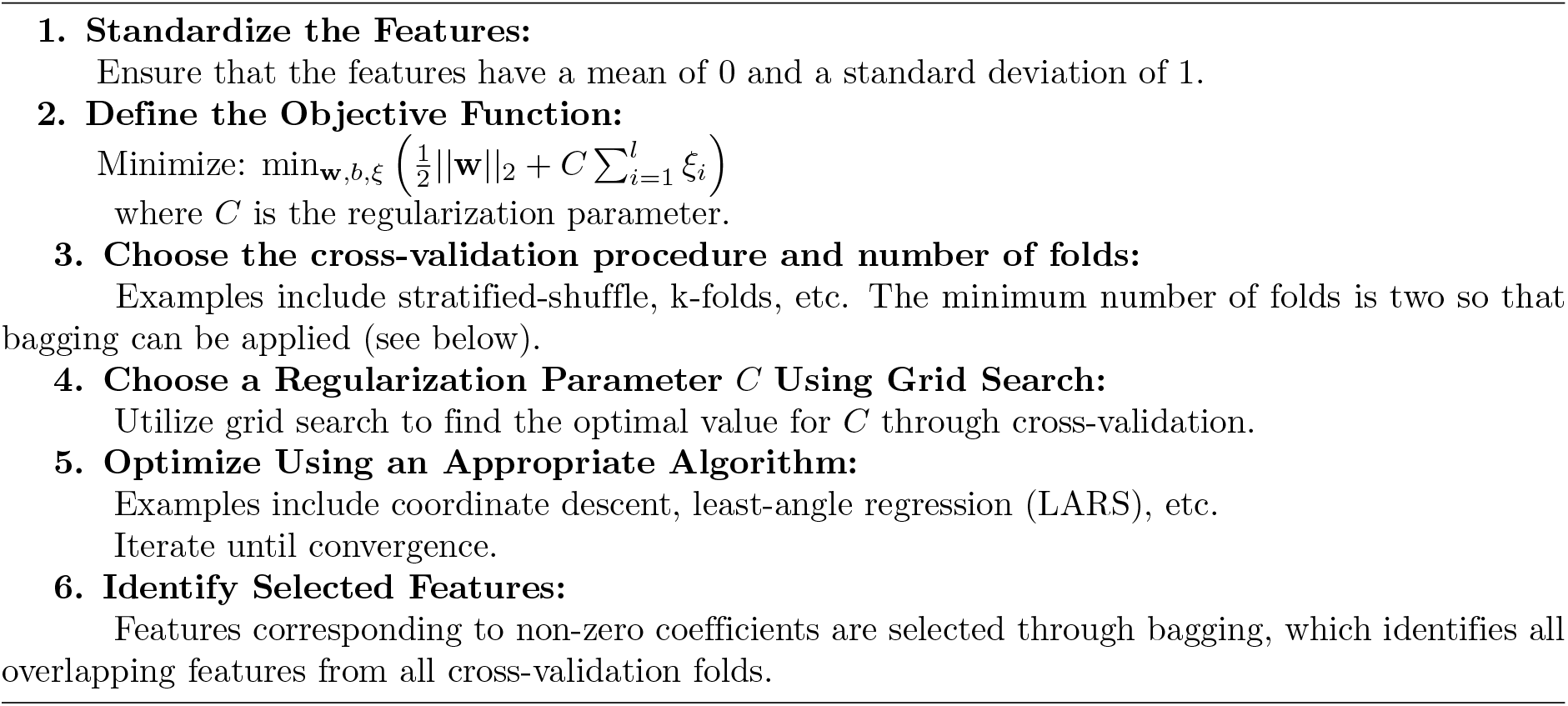

Here, true positives (*TP*) and true negatives (*TN*) represent the number of correctly classified interacting protein pairs. Likewise, false positives (*FP*) and false negatives (*FN*) denote the count of incorrectly classified protein-protein interactions. Accuracy is a suitable measure when the classes in your dataset are well-balanced, meaning there’s roughly an equal number of instances for each class.

Another evaluating metric is *Recall* which measures the proportion of actual positive instances that the model correctly identified,

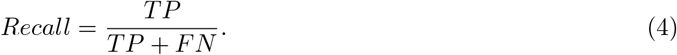

However, *Recall* only looks at the positive class, and sometimes there is a need for a metric that considers both classes.

In addition to *Accuracy* and *Recall*, the model performance can be assessed using a so-called *F*_1_ score (also known as *F* -score or *F* -measure) [15]. It is particularly useful in situations where the data are imbalanced [57], though Matthew’s Correlation Coefficient (*MCC*) has been shown to be more representative with imbalanced datasets [10]. The *F*_1_ score is defined as

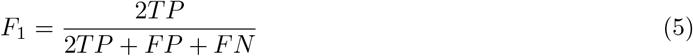

However, one limitation of the *F*_1_ score is that it still doesn’t take true negatives into account. In some cases, correctly identifying negatives (e.g., correctly identifying healthy patients in a medical test) can be just as important as identifying positives.

The final evaluating quantity is a Matthews Correlation Coefficient (*MCC*) that serves as a more dependable statistical measure for complex scenarios,

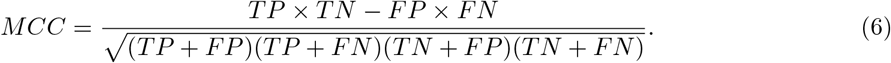

*MCC* ranges from -1 to +1, where +1 represents a perfect prediction, 0 is no better than the random prediction, and -1 indicates total disagreement between prediction and observation.

## 3 Results and Discussions

The results of the feature selection methods for protein-protein interaction networks in EC2, SC5, and SP organisms are shown in Figs. 2, 3, and 4, respectively. In these graphs, negative scores for features indicate that the differences between two proteins’ properties negatively correlate with their ability to interact, while positive scores suggest that those differences positively correlate with the protein-protein interaction.

**Figure 2.**
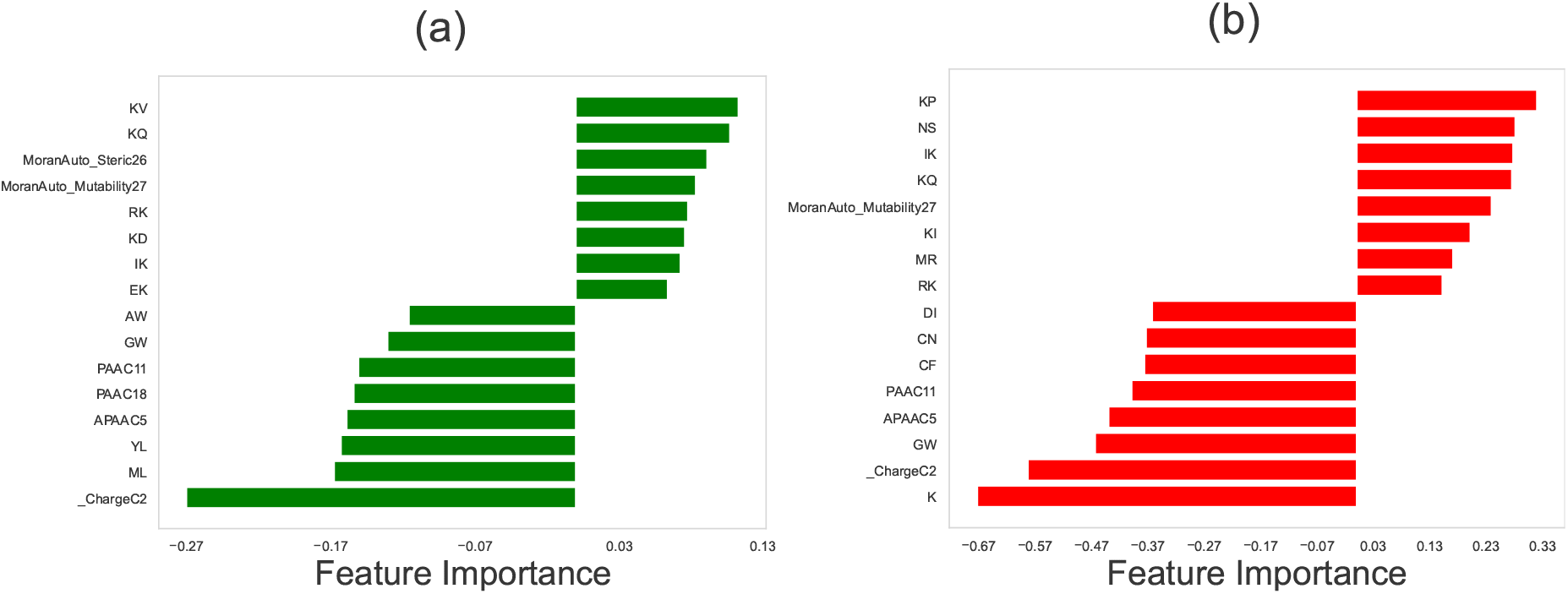
Relative importance of different physicochemical features in identifying the protein-protein interactions in *Escherichia coli* (EC2) network using (a) LASSO regression method, and (b) the Support Vector Machine (SVM). In computations, we utilized the following values for the hyperparameters: for LASSO, the hyperparameter (in Algorithm 1) was set to be *λ* = 0.004. For SVM, the hyperparameter *C* (in Algorithm 2), which is calculated via the grid search optimization, is equal to *C* = 0.1. In both methods, the number of stratified shuffled cross-validation sets is equal to *n* = 18.

**Figure 3.**
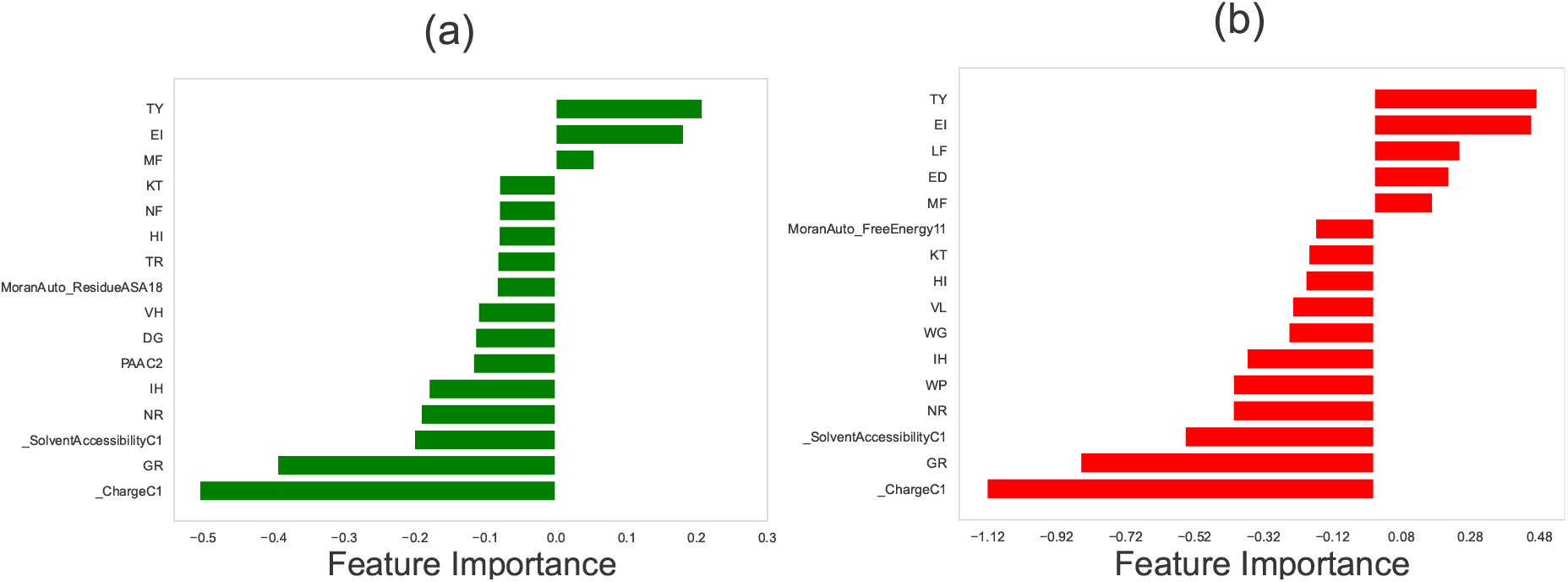
Relative importance of different physicochemical features in identifying the protein-protein interactions in *Saccharomyces cerevisiae* (SC5) network using (a) LASSO regression method, and (b) the Support Vector Machine (SVM). In computations, we utilized the following values for the hyperparameters: for LASSO, the hyperparameter (in Algorithm 1) was set to be *λ* = 0.004. For SVM, the hyperparameter *C* (in Algorithm 2), which is calculated via the grid search optimization, is equal to *C* = 0.1. In both methods, the number of stratified shuffled cross-validation sets is equal to *n* = 15.

**Figure 4.**
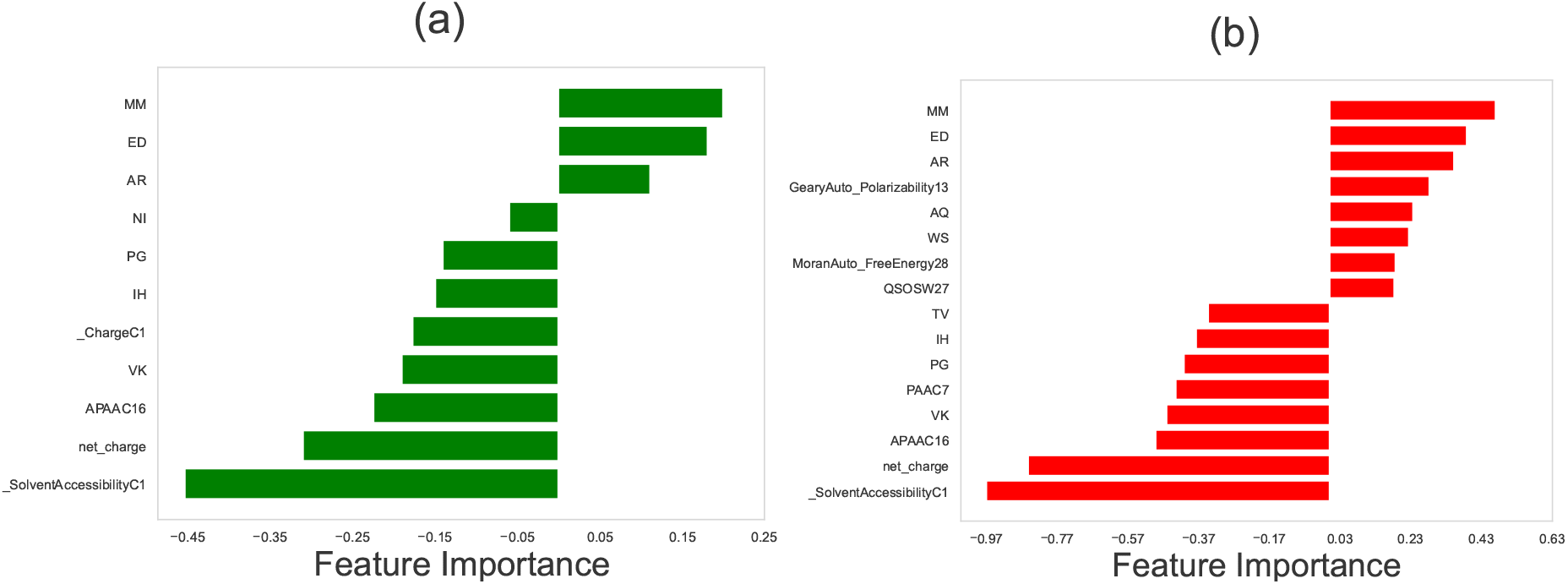
Relative importance of different physicochemical features in identifying the protein-protein interactions in *Schizosaccharomyces pombe* (SP) network using (a) LASSO regression method, and (b) the support vector machine. In computations, we utilized the following values for the hyperparameters: for LASSO, the hyperparameter (in Algorithm 1) was set to be *λ* = 0.004. For SVM, the hyperparameter *C* (in Algorithm 2), which is calculated via the grid search optimization, is equal to *C* = 0.1. In both methods, the number of stratified shuffled cross-validation sets is equal to *n* = 15.

### 3.1 Feature Selection for PPI network in *E. coli* (EC2)

Our feature selection analysis for the protein-protein interactions network in *E. coli* (EC2) has provided interesting insights. Particularly, we observed that differences in dipeptide compositions between two proteins can exhibit both negative and positive effects on their protein-protein interactions (Fig. 2).

Dipeptide composition here represents the fraction of each possible dipeptide (a sequence of two amino acids) within the peptide. Given that there are 20 standard amino acids, there are 20 *×* 20 = 400 possible dipeptides. In the dipeptide composition (DPC), a protein sequence is transformed into a fixed-length feature vector of size 400. Each element of this vector corresponds to one of the possible dipeptides and is calculated as the fraction of the total number of occurrences of that dipeptide in the sequence to the total number of all dipeptides in the sequence. For a protein with *N* amino acids, the total number of dipeptides is *N –* 1, and we have

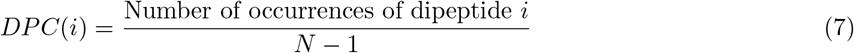

Thus, *DPC*(*i*) for dipeptide *i* is a number between 0 and 1, which corresponds to the probability of finding the dipeptide in the given protein sequence.

The impact of dipeptide compositions on protein-protein interactions (PPI) can be attributed to several reasons. First, the specific arrangement of dipeptides can influence the structural conformations of proteins, affecting their interactions [14]. Second, dipeptide compositions may contain critical binding sites that facilitate or hinder PPI [48, 49]. Third, the presence of charged amino acids in dipeptides can lead to electrostatic interactions that modulate PPI, especially between positively- and negatively-charged amino acids [53]. Fourth, differences in hydrophobicity within dipeptides can also influence interactions, particularly hydrophobic interactions [35].

Our analysis also shows that other selected features such as differences in Amphiphilic Pseudo Amino Acid Composition (APAAC) and Pseudo Amino Acid Composition (PAAC), as predicted by both LASSO and SVM feature selection methods, negatively correlate with protein-protein interactions. These differences may lead to structural incompatibility, altering the distribution of hydrophobic and hydrophilic residues along protein sequences and affecting binding site accessibility [29]. APAAC and PAAC variation might also correspond to hydrophobic-hydrophilic interactions and electrostatic repulsion, reducing the likelihood of stable binding [39, 41]. Moreover, the impact of APAAC and PAAC on PPIs can be context-specific, depending on the organism’s biology and cellular environment. The cumulative effect of these factors can hinder the formation of stable protein complexes and weaken the interactions between proteins, leading to a negative impact on PPI.

Another feature that positively correlates with PPIs in EC2 (see Fig. 2), is differences in Moran’s autocorrelation of mutability and steric properties of the amino acids at certain distances. For example, *MoranAuto Mutability27* refers to Moran’s autocorrelation function of mutability for amino acids that are 27 positions apart in a protein sequence. It means that the mutability in proteins corresponds to the likelihood or rate at which the amino acid residues in a protein sequence change over time due to mutations. This can be influenced by various factors, such as the structural and functional constraints on the protein, as well as the physicochemical properties of the amino acids themselves. The Moran autocorrelation function, which is similar to Pearson’s correlation between the mutability of residue *i* and residue *i* + *d*, is defined as [38]

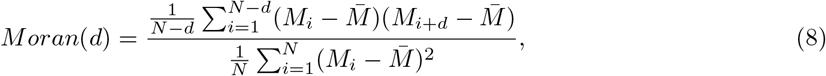

where 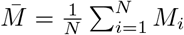 is the average mutability of the sequence. A positive Moran’s value indicates that amino acids that are *d* positions apart in the protein sequence tend to have similar mutability values. While a negative Moran’s value indicates that amino acids that are *d* positions apart in the protein sequence tend to have dissimilar mutability values.

Our feature selection methods suggest that large differences between two proteins in terms of distribution of mutability in the sequences correlate with protein-protein interactions. It could mean that proteins with similar patterns of mutability at a distance of 27 amino acids are more likely to interact with each other. This could potentially be related to how the proteins fold and fit together, as similar patterns of mutability might lead to complementary structural features that facilitate the interaction. Alternatively, it could be related to functional similarities between the interacting proteins, such that they are subject to similar evolutionary pressures that affect their mutability in a coordinated way. Further analysis and validation would be needed to understand the underlying mechanisms behind this association fully.

### 3.2 Feature Selection for PPIs in *S. cerevisiae* (SC5), and *S. pombe* (SP)

For SP and SC5 systems, the protein-protein interaction networks using both LASSO and SVM methods again predict that differences in dipeptide compositions exhibit the strongest correlation with protein-protein interactions: see Figs. 3 and 4. Thus, the role of dipeptide composition in PPIs is not context-specific, indicating that it might be a universal phenomenon valid across all organisms. To test this idea, we applied our analysis to three different multicellular organisms (see the Supporting Information), and it was found that differences in dipeptide compositions also strongly correlate with protein-protein interactions for multicellular organisms, supporting the hypothesis of the universality of dipeptide compositions as a predictor of PPIs.

Moreover, our computational approach predicts that differences in solvent accessibility of two proteins have a negative correlation with PPIs. Solvent accessibility measures how accessible the individual amino acids are to the solvent molecules (typically water) in the protein’s environment [43]. Differences in solvent accessibility between two proteins can have diverse implications on their interactions. Steric hindrance may arise when exposed regions of one protein obstruct the buried regions of the other, hindering effective interaction. Distinct hydrophobic and hydrophilic regions influenced by solvent accessibility may impact the affinity of hydrophobic interactions. Surface complementarity might play a role here: proteins with complementary solvent-accessible surfaces are more likely to form stable interactions. Electrostatic interactions can also be influenced by charged residue exposure, leading to attractive or repulsive forces. Additionally, solvent accessibility may influence conformational changes, affecting the propensity for structural alterations upon interaction. Overall, these factors collectively contribute to the potential impact of solvent accessibility on protein-protein interactions.

### 3.3 Prediction of Protein-Protein Interactions Using Selected Features

After extracting the most important physicochemical properties of each PPI network, our objective is to utilize those features in accurately predicting protein-protein interactions. The performance metrics used for comparison include *Accuracy, Recall*, Matthews Correlation Coefficient (*MCC*), and *F* 1 score as described above. We used SVM and LASSO methods for classifying interacting versus non-interacting protein pairs. As shown in Table 2, selected features from the SVM method generally lead to slightly higher metrics.

**Table 2.**
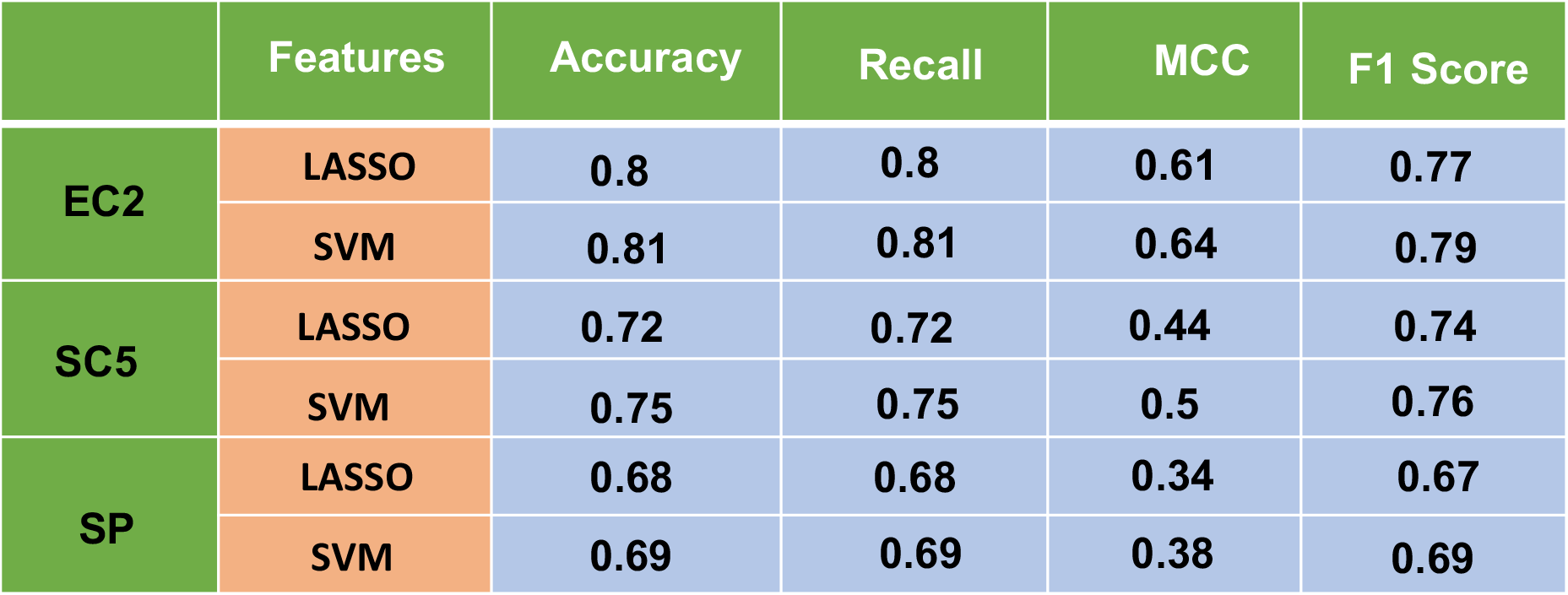
Results of feature selection for protein-protein interactions in *Escherichia coli* (EC2), *Saccharomyces cerevisiae* (SC5), and *Schizosaccharomyces pombe* (SP) networks. Comparison of Accuracy, Recall, Matthews’s correlation coefficient (MCC), and F Score for the trained baseline models (SVM) using selected features from SVM and LASSO. Each metric reflects the average value among 15 test cross-fold validation sets. A standard splitting of 80/20 (training/test) was applied for each fold.

|  | Features | Accuracy | Recall | MCC | F1 Score |
| --- | --- | --- | --- | --- | --- |
| EC2 | LASSO | 0.8 | 0.8 | 0.61 | 0.77 |
|  | SVM | 0.81 | 0.81 | 0.64 | 0.79 |
| SC5 | LASSO | 0.72 | 0.72 | 0.44 | 0.74 |
|  | SVM | 0.75 | 0.75 | 0.5 | 0.76 |
| SP | LASSO | 0.68 | 0.68 | 0.34 | 0.67 |
|  | SVM | 0.69 | 0.69 | 0.38 | 0.69 |

### 3.4 An Illustrative Example of Protein-Protein Interactions: Two-Component PhoB-PhoR System in *E. coli*

To understand better our computational approach, let us apply it for identifying the protein-protein interactions in a two-component system in *E. coli* bacteria. We specifically focus on interactions of histidine kinase PhoR proteins with transcriptional response regulators PhoB proteins [11, 25]. The PhoB-PhoR system in *E. coli* functions to detect low phosphate levels in the environment. When the amount of phosphate species in the medium is low, the PhoR proteins activate the PhoB proteins. The activated (phosphorylated) PhoB proteins then activate genes that help the bacteria absorb more phosphate molecules and use them more efficiently. This system ensures that *E. coli* gets enough phosphate molecules, a vital nutrient, even when they are limited in the surroundings.

Our objective is to show that differences in certain physicochemical features between a PhoR and PhoB correlate with their abilities to interact. We chose an arbitrary response regulator NarL, for which it is known that it does not interact with PhoR. The response regulator NarL is a part of the NarL-NarX/NarQ two-component system. While both protein systems (PhoB-PhoR and NarL-NarX/NarQ) are two-component regulatory systems in *E. coli*, they are tuned to detect and respond to different environmental signals and thus they have distinct regulatory outcomes. In Fig. 5 we compared three proteins in terms of contributions of four dipeptide compositions VV, KK, IK, and EE. One can see that PhoR and PhoB contain different compositions of the corresponding dipeptides, while PhoR and NarL are similar. This suggests that strong differences in the dipeptide compositions correlate with the abilities of these proteins to interact.

**Figure 5.**
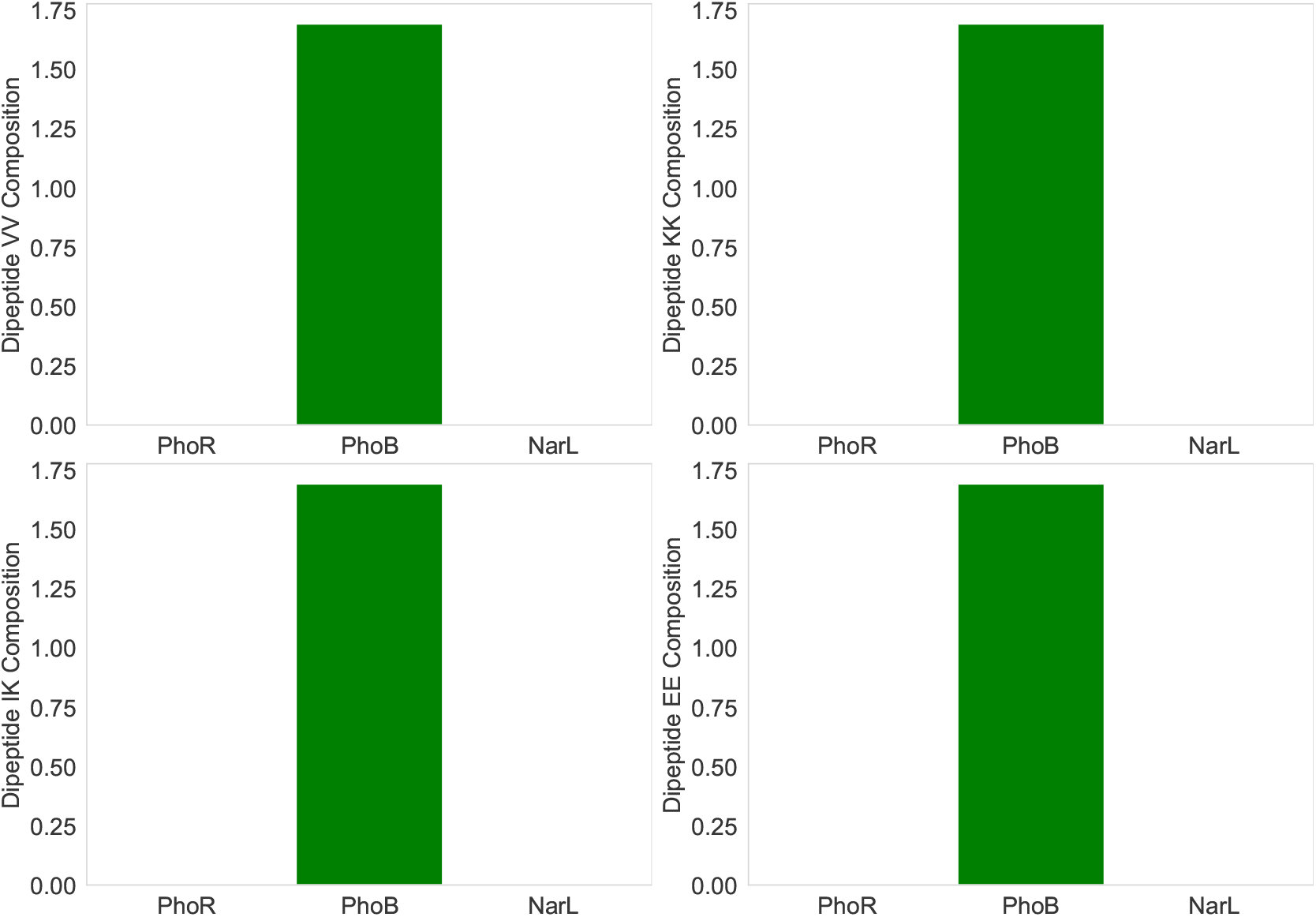
Comparison of charged dipeptide compositions of histidine kinase PhoR with two response regulators PhoB (interacting) and NarL (non-interacting).

The interactions between histidine kinases (HK) and response regulators (RR) in two-component systems (TCS) are governed by specific protein-protein properties that can be significantly affected by dipeptide and amino acid composition differences. The following molecular pictures might be proposed. The specific amino acid sequences and dipeptides at the interaction interfaces of HK and RR determine their cognate pairings, ensuring that a particular HK interacts with its intended RR. Changes in these sequences could disrupt this specificity. The efficiency of phosphoryl transfer between the conserved histidine of HK and the conserved aspartate of RR can be influenced by the surrounding amino acids. For instance, any changes in the nearby residues that might hinder the approach of RR to HK could affect this transfer.

Furthermore, the strength or affinity of the interaction between HK and RR can be controlled by the nature of the amino acids and dipeptides at the binding interface. Hydrophobic, ionic, and hydrogen bond interactions contribute to this binding, and changes in these residues can either enhance or diminish the affinity. HK and RR undergo conformational changes during their interaction. Amino acid or dipeptide composition differences can impact the protein’s ability to undergo necessary conformational changes, which can in turn affect their interaction dynamics. Over evolutionary timescales, if one protein (either HK or RR) changes its amino acid composition and affects its interaction, the interacting partner might co-evolve to accommodate or compensate for this change, maintaining the interaction. This might be the main reason for strong correlations between the differences in the amino-acid and dipeptide compositions and the abilities of proteins to interact.

## 4 Summary and Conclusions

In this study, we introduced a novel computational method of assessing the probability of protein-protein interactions. It is based on the supervised machine learning method for predicting PPIs via correlation analysis by utilizing only protein sequences and their physicochemical properties. A comprehensive set of physicochemical descriptors for proteins is extracted, allowing one to view the properties of protein molecules as a vector in the space of all descriptors. Subsequently, PPIs were studied using a distance vector between pairs of these properties. We performed our analysis on six different datasets belonging to various domains of life: three types of microorganisms (Saccharomyces cerevisiae, Schizosaccharomyces pombe, and Escherichia coli), mammal species (Mus musculus), insects (Drosophila melanogaster), and plants (Arabidopsis thaliana). Despite their differences in complexity, all these organisms rely on PPI networks to perform essential biological functions. These organisms have well-characterized genomes and proteomes, enabling the study of their protein-protein interactions. Examining their PPI networks allows us to gain more microscopic insights into the cellular processes and functions specific to each organism.

We utilized two feature selection methods, LASSO and SVM, to select the most important set of physicochemical descriptors, which have a positive or negative correlation with the protein-protein interactions. These methods reveal that, for all organisms, the differences of two proteins in terms of dipeptide compositions are critically important for identifying PPIs. This is a universal feature that seems to work for all organisms that we investigated. Furthermore, our feature selection methods suggest that there are other physicochemical features specific to each organism that contribute to protein-protein interaction. These types of features, however, are context-dependent [55]. They might be specific to the organism’s biology, the cellular environment, or the specific protein network being considered. Different organisms or cell types might have distinct requirements for protein interactions, leading to different preferences for certain physicochemical properties [34, 47].

The impact of correlations of dipeptide compositions with PPIs can be attributed to several sources. First, it could be related to the structural conformations of proteins. Dipeptides are short sequences of two amino acids, and their specific arrangement can influence the overall secondary and tertiary structure of proteins [27]. The three-dimensional structure of proteins is important in determining how they interact with other proteins [28]. Differences in dipeptide compositions might also lead to the variations in protein folding [16, 40], which, in turn, can affect their ability to interact with other proteins. Second, dipeptide compositions may contain specific amino acid pairs that serve as critical binding sites for PPIs. These binding sites can mediate physical interactions between proteins and are essential for the formation of protein complexes. Variations in dipeptide compositions can alter the presence or accessibility of these binding sites, influencing the potential for PPIs. Third, amino acids in dipeptide compositions can have different charges, such as positively charged (e.g., lysine), negatively charged (e.g., aspartic acid), or neutral (e.g., alanine). These charged amino acids can engage in electrostatic interactions with other proteins, either promoting or inhibiting their interactions. Dipeptides with specific combinations of charged amino acids may create favorable or unfavorable electrostatic environments for PPI. Finally, some dipeptide compositions may contain hydrophobic amino acids, which tend to cluster together in the protein’s core [3, 51], while others may have hydrophilic amino acids exposed on the protein’s surface [1, 46]. Differences in dipeptide compositions can lead to variations in hydrophobic and hydrophilic regions, influencing protein-protein interactions, especially those that involve hydrophobic interactions, and dipeptide composition can be targeted to affect protein-protein interactions [26].

Understanding the physicochemical properties of the interface formed by protein-protein association might help to clarify the mechanisms of formation of protein interaction networks on one hand, and to design molecules that can engage with a given interface and thereby control protein function on the other hand. For example, synthetic molecules that resemble the chemical structure of proteins, called peptidomimetics, can be used to inhibit protein-protein interactions associated with diseases [18]. This means that PPIs might be excellent targets for drug development [2]. By considering the specific physicochemical features of each PPI network, our computational approach can capture the network-specific patterns and relationships that govern protein-protein interactions in different biological contexts. This allows for a more accurate and context-specific prediction of protein-protein interactions, enhancing our understanding of how these interactions contribute to cellular processes and functions in each organism.

It’s important to note that protein-protein interactions are highly complex and multifaceted processes, involving various molecular forces and structural features. Machine learning models can help to identify patterns and correlations in large datasets, but they may not capture the full microscopic intricacies of protein-protein interactions. As with any predictive model, it is essential to interpret the results cautiously and complement them with experimental validations and further analysis to gain a deeper understanding of the underlying biology. Additionally, considering other physicochemical properties and features in combination with solvent accessibility can lead to a more comprehensive understanding of protein-protein interactions. Our method can be applied to PPI systems in humans, including virus–host systems [4] and cancer [23, 56].

## Data and Software Availability

The data obtained in this work and the in-house scripts are available on GitHub at the following URL: https://github.com/hamid-teimouri/PPI_dipeptide_similarity.git

## Author contributions

H.T. and A.M. designed the research. H.T. and A.M. performed the research. H.T., A.M., and A.B.K. wrote the article. All authors reviewed the article.

