## Supporting Information for "Physical-Chemical Features Selection Reveals That Differences in Dipeptide Compositions Correlate Most with Protein-Protein Interactions"

In this supporting information we present additional figures and tables that were referenced in the main text.

### Feature selection in determining the protein-protein interactions in multicellular organisms

Results of feature selection for three types of multicellular organisms, including *Arabidopsis thaliana* (AT), *Mus musculus* (MM), and *Drosophila melanogaster* (DM2) are presented in figures S1, S2, and S3, respectively.

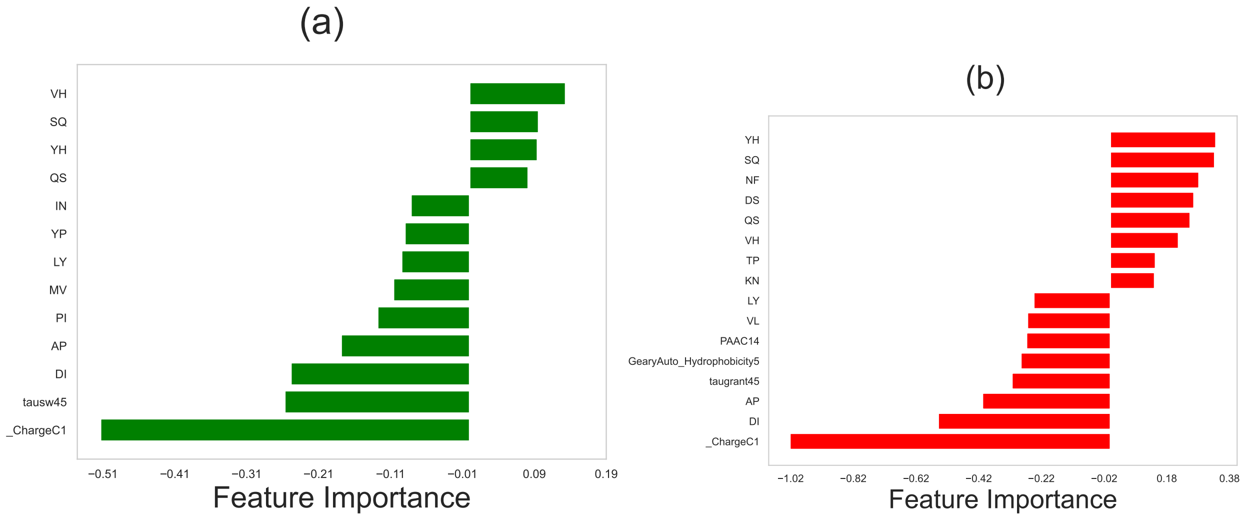


Figure S1: Relative importance of different physicochemical features in determining the protein-protein interactions in *Arabidopsis thaliana* (AT) network using (a) LASSO regression method, and (b) the support vector machine. In computations, we utilized the following values for the hyperparameters: For LASSO, the hyperparameter was set to be $\lambda = 0.004$. For SVM, the hyperparameter $C$ , which is calculated via the grid search optimization, is equal to $C = 0.1$. In both methods, the number of stratified shuffled cross-validation sets is $n = 15$.

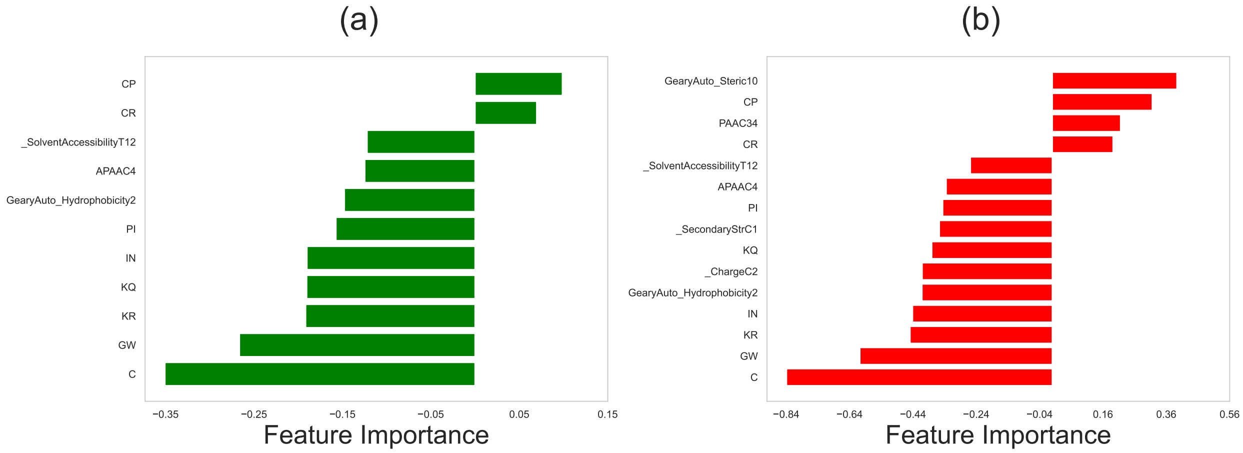


Figure S2: Relative importance of different physicochemical features in determining the protein-protein interactions in *Mus musculus* (MM) network using (a) LASSO regression method, and (b) the support vector machine. In computations, we utilized the following values for the hyperparameters: For LASSO, the hyperparameter was set to be $\lambda = 0.004$. For SVM, the hyperparameter $C$ , which is calculated via the grid search optimization, is equal to $C = 0.1$. In both methods, the number of stratified shuffled cross-validation sets is $n = 15$.

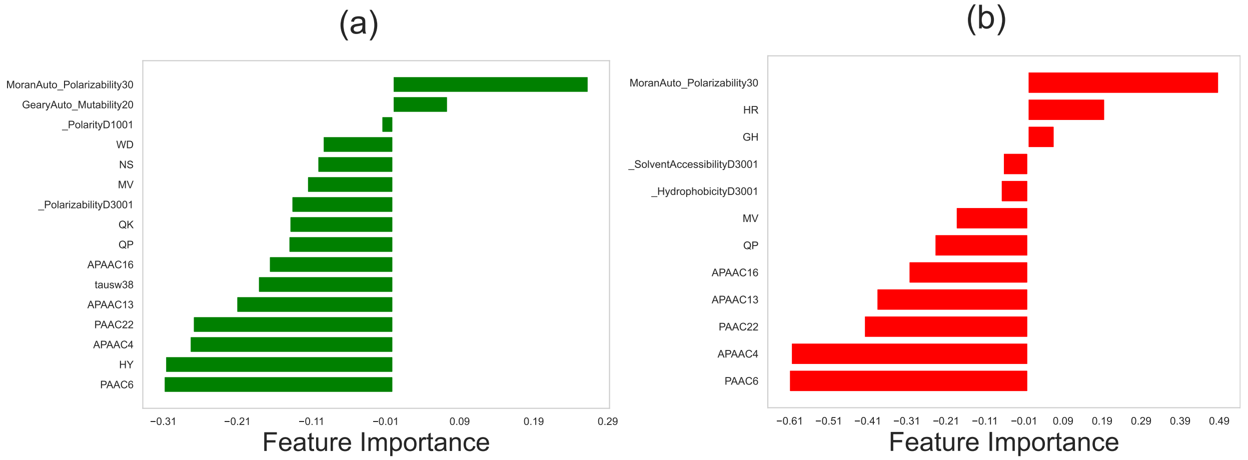


Figure S3: Relative importance of different physicochemical features in determining the protein-protein interactions in *Drosophila melanogaster* (DM2) network using (a) LASSO regression method, and (b) the support vector machine. In computations, we utilized the following values for the hyperparameters: For LASSO, the hyperparameter was set to be $\lambda = 0.004$. For SVM, the hyperparameter $C$ , which is calculated via the grid search optimization, is equal to $C = 0.1$. In both methods, the number of stratified shuffled cross-validation sets is $n = 15$.

### Model evaluation for prediction of protein-protein interactions in multicellular organisms

Results of support vector machine and LASSO methods (using selected features) for classifying interacting from non-interacting protein pairs for PPI networks in *Arabidopsis thaliana* (AT), *Mus musculus* (MM), and *Drosophila melanogaster* (DM2) are presented in Table S1.

| | Features | Accuracy | Recall | MCC | F1 Score |
|---|---|---|---|---|---|
| AT | LASSO | 0.66 | 0.66 | 0.33 | 0.67 |
| | SVM | 0.69 | 0.69 | 0.38 | 0.69 |
| MM | LASSO | 0.66 | 0.66 | 0.31 | 0.64 |
| | SVM | 0.69 | 0.69 | 0.38 | 0.68 |
| DM2 | LASSO | 0.72 | 0.72 | 0.45 | 0.72 |
| | SVM | 0.71 | 0.71 | 0.42 | 0.71 |

Table S1: Results of feature selection for protein-protein interactions in *Arabidopsis thaliana* (AT), *Mus musculus* (MM), and *Drosophila melanogaster* (DM2) networks. Comparison of Accuracy, Recall, Matthews's correlation coefficient (MCC), and F1 Score for the trained baseline models (SVM) using all features vs the models trained using selected features from SVM and LASSO. Each metric reflects the average value among 15 test cross-fold validation sets. A standard splitting of 80/20 (training/test) was applied for each fold.
